# A TaqMan Assay That Knows Its Mite: Specific Detection of *Tropilaelaps mercedesae* Clade I

**DOI:** 10.1101/2025.11.07.686760

**Authors:** Marga van Gent-Pelzer, Marc Hendriks, Terd Disayathanoowat, Patcharin Phokasem, Heather Graham, Delphine Panziera

## Abstract

*Tropilaelaps mercedesae* (*Mesostigmata*: *Laelapidae*), an invasive mite native to Southeast Asia, is spreading rapidly due to global bee trade and climate change. Its detection in Eastern Europe raises concerns about its entry into the EU. As a notifiable parasite under EU Animal Health Law, early and reliable detection is essential. We developed a sensitive TaqMan assay targeting the cytochrome C oxidase subunit I (COI) gene, using synthetic double-stranded DNA fragments (gBlocks). This assay enables rapid, specific, and scalable detection of *T. mercedesae* Clade I.

---

To support assay development, *Tropilaelaps* spp. *(Mesostigmata: Laelapidae)* mites from *Apis mellifera* colonies in Chiang Mai province were collected by hand and stored in RNAlater (Invitrogen by Thermo Fisher Scientific, Vilnius, Lithuania). DNA was extracted from individual mites using a disposable pestle, grinding the dried off specimens in microtubes. The crushed mites were then manually homogenized in lysis buffer either from the RNeasy Micro Kit (A) or the DNeasy Blood & Tissue Kit (B) (Qiagen Benelux B.V., Venlo, the Netherlands) following the manufacturer’s protocols. Two additional kits (C and D); were evaluated for their suitability (see Supplementary Figure 1).

**Figure 1.**
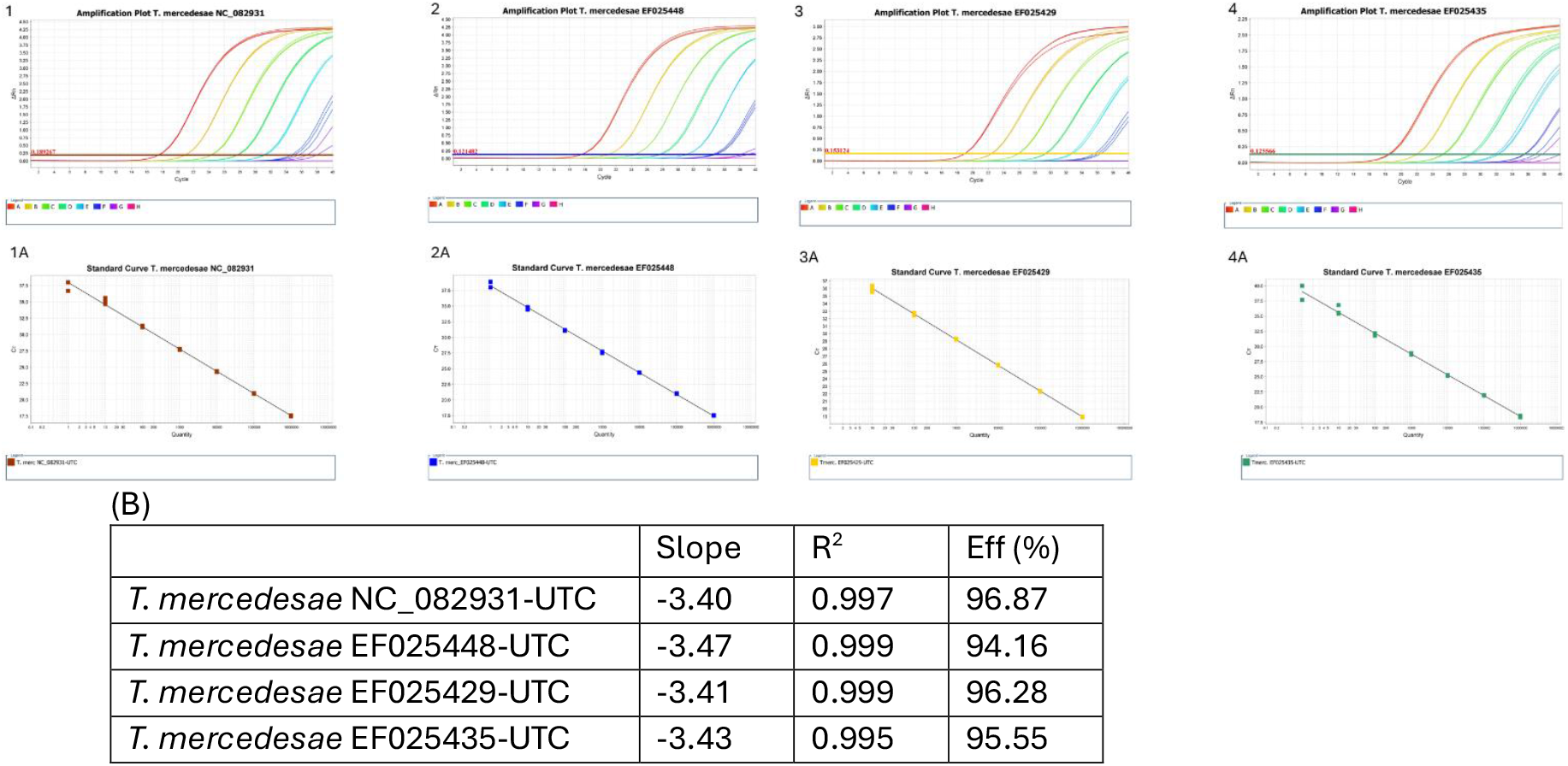
Amplification plots and standard curves of the *Tropilaelaps mercedesae* (Clade I) TaqMan assay. NC_082931 (1 and 1A), EF025448 (2 and 2A), EF025429 (3 and 3A), EF025435 (4 and 4A). Ct values of gBlock constructions serial dilutions were plotted against the log value of the target DNA amount (log conc). (B) Slope, the coefficient of correlation (R^2^) and efficiency of the reaction (Eff%) represents the result of a triplicate amplification of each dilution.

The species identity of the collected mites was confirmed by amplicon sequencing, two primer sets, LCO1490 and HCO2198 (Folmer *et al*, 1994) and TCF1 and TCR2 (Anderson and Morgan, 2007) amplifying overlapping regions of the cytochrome C oxidase subunit I gene (COI) were used for PCR amplification. PCR reactions were carried out in a total volume of 25 µl containing repliQa HiFi ToughMix (Quantabio, Beverly, MA, USA), 300 nM of each primer, 2 µl of DNA template, and nuclease-free water. Thermal cycling conditions consisted of 40 cycles of 98°C for 10 seconds, followed by 5 seconds at 55°C and 5 seconds at 68°C.

PCR products were sequenced at Macrogen Europe B.V. (Amsterdam, the Netherlands). COI sequences from five Thai specimens, each 828 base pairs in length, were assembled using CLC Genomics Workbench (Qiagen Denmark, version 24.0.2) and deposited in the GenBank database under accession numbers PX125996–PX126000. The obtained sequences were compared to *Tropilaelaps* entries in the NCBI GenBank database using BLAST (ref) and the best match was with the COI gene specific to *T. mercedesae*. Subsequently, alignments were performed with NC_082931 as the reference COI gene. All available accessions up to January 2025 were included in the analysis. *In silico* screening and data analysis identified a conserved 211-bp region with high sequence similarity across *T. mercedesae* entries.

A TaqMan assay was designed using the PrimerQuest tool from IDT (Integrated DNA Technologies, Leuven, Belgium), targeting this region with the following oligonucleotides: Tmer_COI-F1 GCAGCAGCTATTACCATACTTCTA, Tmer_COI-R1 TGTAAACTTCAGGGTGACCAAA and Tmer_COI-P1 FAM-CAGGTGGAG-ZEN-GAGACCCAATCCTTT-BHQ (IDT). To validate the assay, synthetic double-stranded DNA fragments (gBlocks) representing target sequences were used as positive and negative controls, mimicking real-world DNA samples and confirming assay specificity (Conte et al., 2018).

Based on the Maximum Likelihood phylogenetic tree described by Namin et al. (2024), gBlocks were designed to represent the various *Tropilaelaps* subclades. Sequences were selected to maximize variation in the selected primers/probe binding sites within *T. mercedesae*, while minimizing similarity to non-target *Tropilaelaps* species. In table 1 the sequence of the targeted *T. mercedesae* accessions can be found: NC_082931 (perfect match, Malay subclade), EF025448 (Mainland), EF025429 (Tibet), EF025435 (Indonesian), EF025440 and EF025442 (Clade II). Also in table 1 the sequence of closest related non-targets *T. clareae* (EF025464, EF025458), *T. thaii* (EF025452), *T. koenigerum* (EF025449) and *Pneumolaelaps fuscicolens* (MW367916) were included.

**Table 1.**
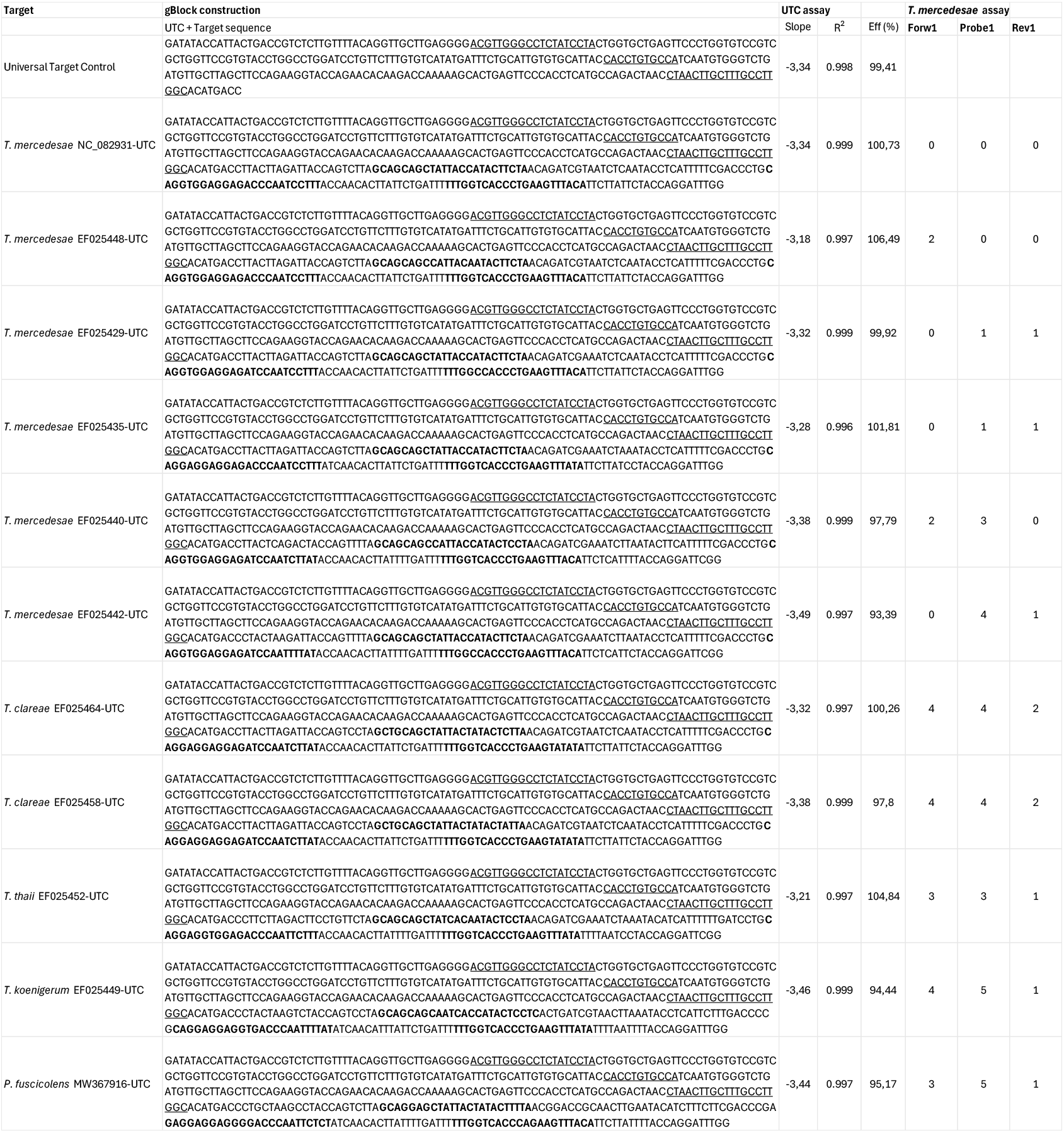
Target information used for TaqMan development based on Universal Target Control (UTC), *Tropilaelaps mercedesae* (Clade I): NC_082931, EF025448, EF025429, EF025435. *T. mercedesae* (Clade II) EF025440 and EF025442. *T. clareae* EF025464 and EF025458, *T. thaii* EF025452, *T. koenigerum* EF025449 and *Pneumolaelaps fuscicolens* MW367916. The primers and probe sequences are indicated in the gBlock construction (underlined in UTC and bold in *Tropilaelaps* spp. and *P. fuscicolens*. The performance of the UTC assay is indicated by Slope, R^2^ and Efficiency. Single Nucleotide Polymorphism’s in primers and probe site for *T. mercedesae* assay are expressed in number.

Each gBlock construction contains the target gene (170 bp) and the Universal Target Control (UTC) (289 bp). The UTC assay serves as a reference to monitor the performance and quality of the gBlock constructions (van Gent-Pelzer et al., 2024), because both targets are present in a 1:1 ratio. The amount of the gBlock constructions was calculated to the copy number and the gBlock constructions were serially diluted 10-fold in TE buffer from 1.0E+06 to 1 copy per µl to create a qPCR standard curve.

TaqMan reactions were performed on the Quantstudio 12 K Flex (Thermo Fisher Scientific, Waltham (MA), USA). Reactions were run in a 25 µl volume consisting of PerfeCta qPCR ToughMix Low ROX (Quantabio, Beverly MA, USA), primers (300 nM) and probe (100 nM), 5 µl DNA or 1 µl gBlock construction. Thermal cycling conditions consisted of 1 min 95 °C followed by 40 cycles of 15 sec at 95 °C and 1 min at 60 °C. No-Template-Control (NTC) reactions were included. Threshold and baseline settings for each run were automatically set by the software. The real-time PCR software v 3.1 provides a standard curve and the amplification efficiency of each TaqMan assay was determined based on the slope of the log-linear portion of the standard curve (E (Efficiency) = [10(-1/slope)]-1). The performance of the twelve gBlock constructions was checked by the amplification efficiency of the UTC assay. The gBlock containing only the UTC sequence, gave a linear curve, with correlation coefficient R^2^ =0.998, slope = −3.34 and efficiency of amplification E =99.41 %. The UTC from the eleven constructed gBlocks generated similar linear standard curves and met the requirements (see Table 1).

The *T. mercedesae* (Clade I) TaqMan assay (Fig. 1) demonstrates high sensitivity, with a limit of detection (LOD) between 1–10 DNA copies. It is also specific and reliably amplifying only the intended target without generating false positives, even when tested against a gBlock construction containing 10^6^ copies of the non-target sequences in the reaction tube. Despite intraspecific variation—such as two SNPs in the forward primer binding site (EF025448-UTC) or one SNP in the probe and reverse primer region (EF025429-UTC and EF025435-UTC)—the assay maintains consistent in analytical performance. Parameters including slope, R^2^, and efficiency are comparable to those obtained with perfectly matched primers and probes (NC_082931-UTC). The gBlock construction *T. mercedesae*_EF025440-UTC (Clade II) does produce a standard curve from Ct values (not shown); however, due to three SNPs in the probe region, the fluorescent signal is significantly reduced (dRn ≈ 0.2), rendering it unsuitable for reliable quantification.

DNA extracted from sixteen individual *T. mercedesae* mites from Thailand was successfully detected and quantified using the *T. mercedesae* (Clade I) TaqMan assay. To test a-specificity, the parasitic mite *Varroa destructor* present in beehives was included. No a-specific signal was found for *Varroa destructor* collected in Thailand and the Netherlands. The 18S TaqMan assay (Silacci et al., 2018) confirmed the high DNA quality in these non-target samples (see Supplementary Figure 2).

These findings confirm that the assay reliably detects *T. mercedesae* (Clade I) and is suitable for detection of a single mite, offering a sensitive and field-applicable tool for early detection and monitoring. As new diversity within the *Tropilaelaps* spp. might be discovered, regular updating the assay when new information becomes available in public databases is recommended to re-validate the specificity and selectivity of the described assays. Sharing collected sequence data from additional *Tropilaelaps* samples, will benefit the development and in silico evaluation of the TaqMan assays.

---

**Supplementary figure S1.**
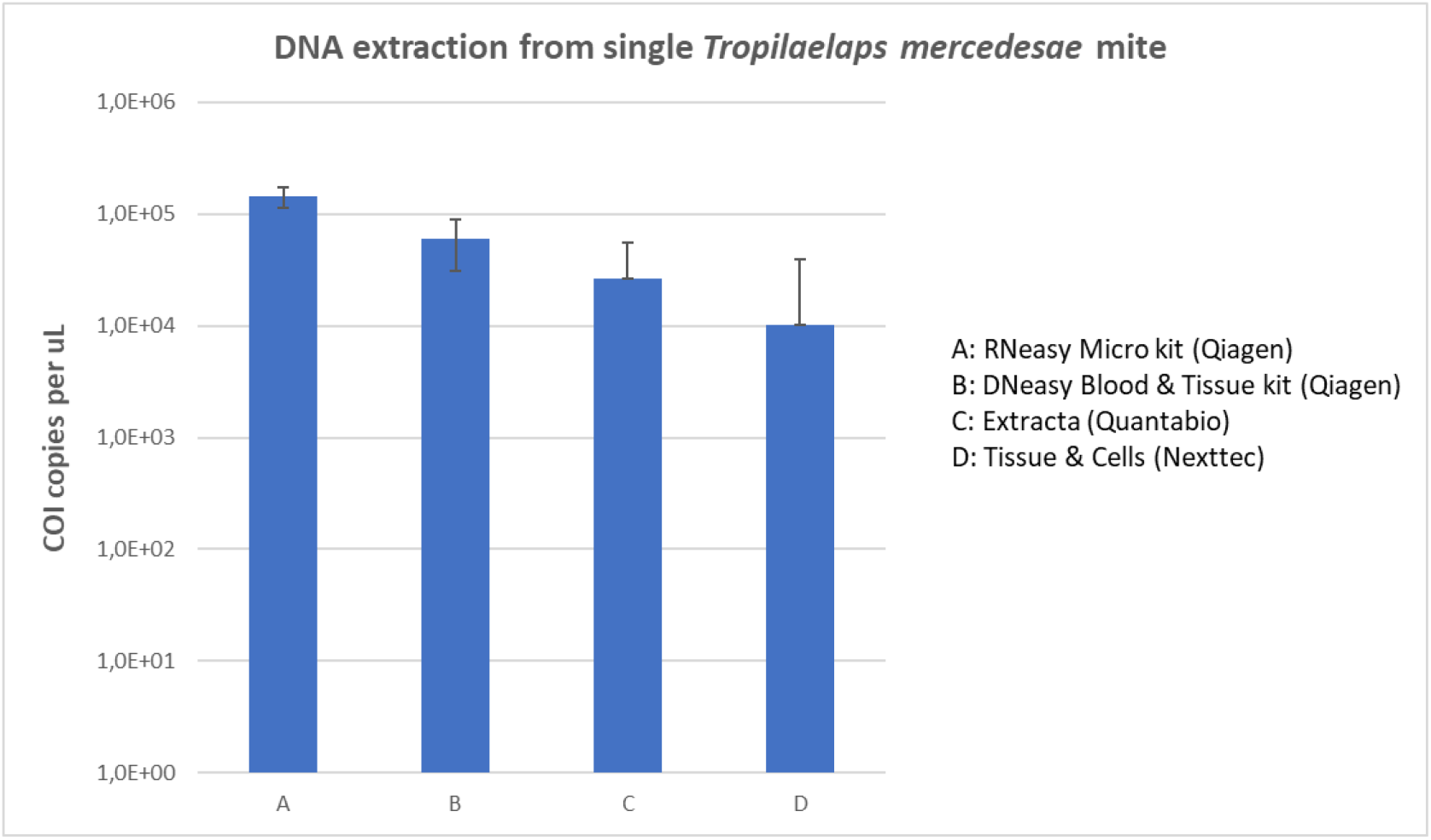
Comparison of four extraction kits (A,B,C,D) for their suitability for DNA extraction from single *Tropilaelaps* mites (n = 4) and TaqMan amplification in COI copies per µl.

**Supplementary figure S2.**
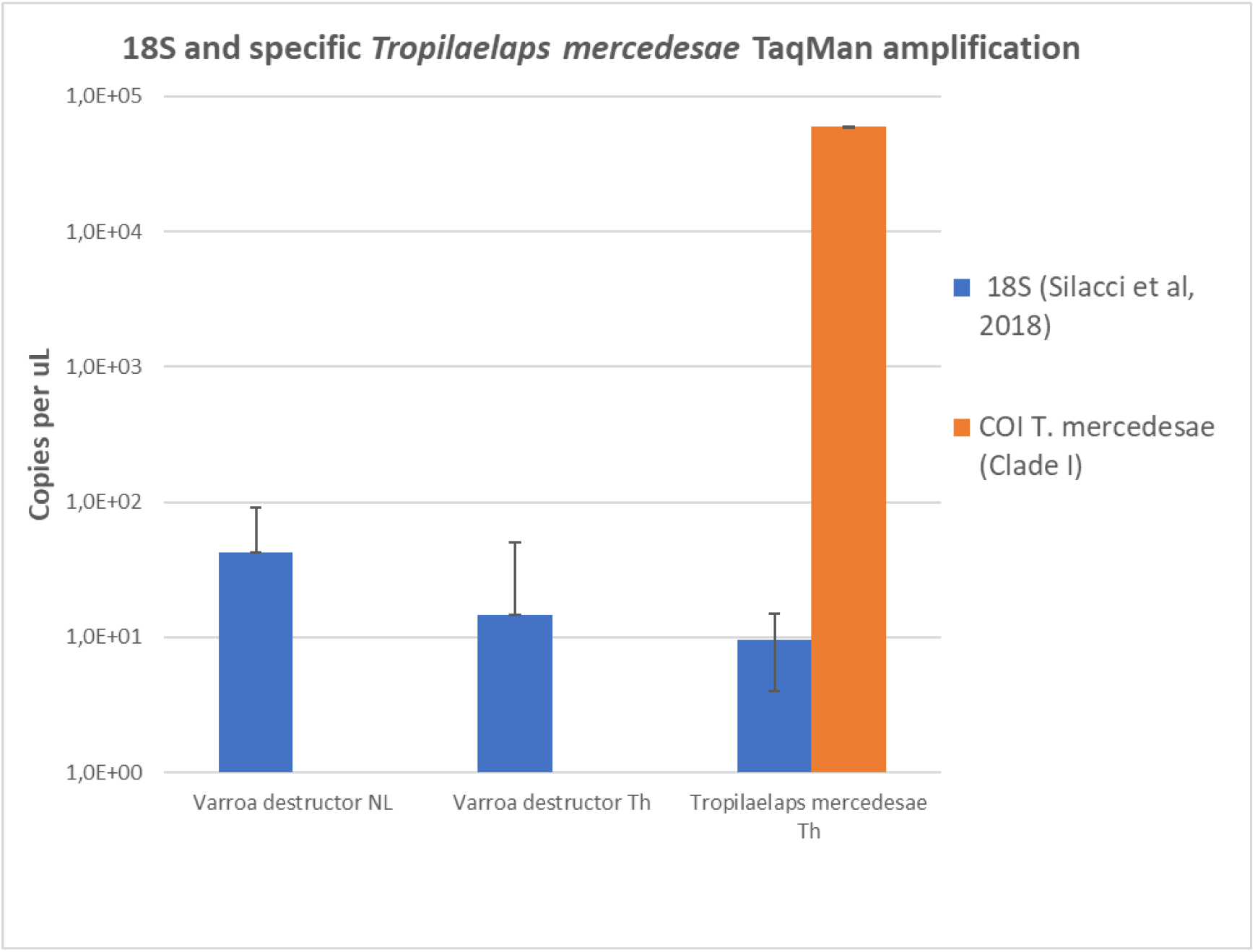
Detection of *Tropilaelaps mercedesae* DNA using the developed species-specific Clade I TaqMan. DNA from individual mites (n=16) collected in the Chiang Mai province in Thailand was successfully amplified. No a-specific signal was observed from *Varroa destructor* mites (n=4) collected in Thailand and the Netherlands. The 18S TaqMan assay (Silacci et al., 2018) confirmed the DNA quality of these samples.

